# *Campylobacter jejuni* modulates reactive oxygen species production and NADPH oxidase 1 expression in human intestinal epithelial cells

**DOI:** 10.1101/2022.03.04.482506

**Authors:** Geunhye Hong, Cadi Davies, Zahra Omole, Janie Liaw, Anna D. Grabowska, Barbara Canonico, Nicolae Corcionivoshi, Brendan Wren, Nick Dorrell, Abdi Elmi, Ozan Gundogdu

**Affiliations:** Faculty of Infectious and Tropical Diseases, London School of Hygiene & Tropical Medicine, Keppel Street, London, WC1E 7HT, UK; Department of Biophysics and Human Physiology, Medical University of Warsaw, Warsaw, 02-091, Poland; Department of Biomolecular Sciences, University of Urbino Carlo Bo, Urbino, 61029, Italy; Bacteriology Branch, Veterinary Sciences Division, Agri-Food and Biosciences Institute (AFBI), Belfast, BT9 5PX, UK

**Keywords:** *Campylobacter jejuni*, Intestinal epithelial cells, NADPH oxidase 1, Reactive oxygen species

## Abstract

*Campylobacter jejuni* is the major bacterial cause of foodborne gastroenteritis worldwide. Mechanistically, how this pathogen interacts with intrinsic defence machinery of human intestinal epithelial cells (IECs) remains elusive. To address this, we investigated how *C. jejuni* counteracts the intracellular and extracellular reactive oxygen species (ROS) in IECs. Our work shows that *C. jejuni* differentially regulates intracellular and extracellular ROS production in human T84 and Caco-2 cells. *C. jejuni* downregulates the transcription and translation of Nicotinamide adenine dinucleotide phosphate (NAPDH) oxidase (Nox1), a key ROS-generating enzyme in IECs and antioxidant defence genes *cat* and *sod1*. Furthermore, inhibition of Nox1 by diphenylene iodonium (DPI) and siRNA reduced *C. jejuni* ability to interact, invade and intracellularly survive within T84 and Caco-2 cells. Collectively, these findings provide mechanistic insight into how *C. jejuni* modulates the IEC defence machinery.

## Introduction

Microbial pathogens have evolved to possess subversion strategies to alter the functionality of host cells upon infection (Escoll, Mondino, Rolando, & Buchrieser, 2016). These include modulation of host cell functions that involve vesicle trafficking, apoptosis, and immune activation (Asrat, de Jesús, Hempstead, Ramabhadran, & Isberg, 2014; Pedron et al., 2007; Rudel, Kepp, & Kozjak-Pavlovic, 2010). Crucially, these host cell functions are essential for elimination of foreign pathogens. The evolving battle between pathogen and host adds to the complexity of the pathogenesis of infection (Escoll et al., 2016).

*Campylobacter jejuni* is the leading foodborne bacterial cause of human gastroenteritis worldwide (Silva et al., 2011). *C. jejuni* causes watery or bloody diarrhoea, abdominal pain, and fever. *C. jejuni* infection can also lead to Guillain-Barré Syndrome (GBS), a rare but severe post-infectious autoimmune complication of the peripheral nervous system (Kaakoush, Castaño-Rodríguez, Mitchell, & Man, 2015; Silva et al., 2011; Willison, Jacobs, & van Doorn, 2016). Importantly, Campylobacteriosis in low-income countries is associated with child growth impairment and can be fatal in children (Amour et al., 2016). Although *C. jejuni* is a microaerophilic bacterium, its omnipresence in the environment and various hosts is mitigated by regulatory mechanisms against oxidative stress (Gundogdu et al., 2016). Upon adhering and invading human intestinal epithelial cells (IECs), *C. jejuni* manipulates host cytoskeleton regulation to maximise its invasion (Negretti et al., 2021). Following invasion, *C. jejuni* resides in cytoplasmic vacuoles named Campylobacter containing vacuoles (CCVs) which can escape the canonical endocytic pathway and avoid fusion with lysosomes (M. E. Konkel, Hayes, Joens, & Cieplak Jr, 1992; Watson & Galán, 2008). These findings demonstrate that modulation and invasion of host IECs are a prerequisite for human intestinal disease caused by *C. jejuni*.

A vital mechanism used by host cells in response to pathogens is the production of reactive oxygen species (ROS) which are highly reactive molecules, such as oxygen radicals and non-radicals, produced by the partial reduction of oxygen (Aviello & Knaus, 2017). When phagocytes such as macrophages detect and engulf pathogens using the respiratory burst, ROS are rapidly generated to eradicate the engulfed pathogens through oxidative damage (Paiva & Bozza, 2014). Interestingly, the level of ROS produced by human IECs is lower in comparison to resident macrophages and blood leukocytes (neutrophils and monocytes), however, ROS in IECs can also exhibit antimicrobial activity by inducing inflammation (Burgueño et al., 2019; Holmström & Finkel, 2014; Paiva & Bozza, 2014). The precarious nature of ROS production by IECs is demonstrated by exhibiting both deleterious and beneficial host effects, thus homeostasis of ROS is essential. To counter the damaging effects of ROS, host IECs possess antioxidant components that neutralise ROS, such as catalase, superoxide dismutase and glutathione peroxidase. Nicotinamide adenine dinucleotide phosphate oxidase (NADPH oxidase; Nox) and mitochondria have central roles as predominant sources of ROS in human IECS (Aviello & Knaus, 2017).

Nox is an essential multicomponent enzyme which catalyses production of superoxide (O_2_^−^) (Brandes, Weissmann, & Schröder, 2014; Sumimoto, Miyano, & Takeya, 2005). In IECs, the most abundant types of Nox are Nox1 and Nox4. Intriguingly, Nox4 is constitutively active, whereas Nox1 is not. The Nox1 complex is composed of Nox1, p22phox, Nox organiser 1 (NoxO1), Nox activator (NoxA1) and small GTPase Rac1. Nox1 is the catalytic subunit of the complex on the plasma membrane and its activation is dependent on supplementary cytosolic subunits. Following this, p22phox is transported to the plasma membrane promoted by Nox1 expression (Brandes et al., 2014). Upon activation, NoxO1 binds to both NoxA1 and p22phox targeting NoxA1 to the plasma membrane. In turn NoxA1 binds to GTP (guanosine triphosphate)-bound Rac1 and promotes electron flow through flavocytochrome in Nox1 in a GTP-dependent manner. Studies have shown that GTP-bound Rac1 is essential for activity of Nox1 (Nisimoto et al., 2008; Ueyama, Geiszt, & Leto, 2006). Electrons travel from NADPH initially to flavin adenine dinucleotide (FAD), then through the Nox heme groups and finally to oxygen, forming O_2_^−^ (Nisimoto et al., 2008). Notably, Nox1-mediated ROS play important roles in IECs including regulation of growth and proliferation, epithelial wound healing, intestinal host defence, and maintenance of bacterial homeostasis in the GI tract (Juhasz et al., 2017; Lipinski et al., 2019; Matziouridou et al., 2018).

How *C. jejuni* interacts with the inherent defence machinery of human IECs remains unclear. To explore this further, we examined the mechanisms *C. jejuni* uses to counteract the intracellular and extracellular ROS in IECs. Previous findings demonstrated the upregulation of Nox1 in IECs by enteric pathogens such as *Escherichia coli* (Elatrech et al., 2015), *Salmonella* Enteritidis (Kawahara et al., 2016), and *Helicobacter pylori* (den Hartog et al., 2016; Kawahara et al., 2005). However, given the invasion and survival properties of *C. jejuni*, we hypothesised that *C. jejuni* may have distinct host cell modulation mechanisms in play. In this study, we show that diverse *C. jejuni* strains downregulate both intracellular and extracellular ROS production in human IECs by modulating the expression of Nox1. We demonstrate inhibition of Nox1 by diphenylene iodonium (DPI) and siRNA reduced the ability of *C. jejuni* to interact, invade and intracellularly survive within T84 and Caco-2 cells. Our results highlight a unique strategy of *C. jejuni* survival and emphasise the importance of Nox1 in *C. jejuni*-IEC interactions. This represents a distinctive mechanism that *C. jejuni* uses to modulate IEC defence machinery.

## Experimental procedures

### Bacterial strains and growth conditions

*C. jejuni* wild-type strains used in this study are listed in Table S1. For general growth, all *C. jejuni* strains were grown on Columbia Blood Agar (CBA) plates (Oxoid, U.K) supplemented with 7% (v/v) horse blood (TCS Microbiology, UK) and Campylobacter selective supplement Skirrow (Oxoid) at 37°C under microaerobic conditions (10% CO_2_, 5% O_2_ and 85% N_2_) (Don Whitley Scientific, U.K).

### Human intestinal epithelial cell culture

T84 cells (ECACC 88021101) and Caco-2 cells (ECACC 86010202) were obtained from European Collection of Authenticated Cell Cultures (ECACC). T84 and Caco-2 cells were cultured in a 1:1 mixture of Dulbecco’s modified Eagle’s medium and Ham’s F-12 medium (DMEM/F-12; Thermo Fisher Scientific, U.S.A) with 10% Fetal Bovine Serum (FBS; Labtech, U.K), 1% non-essential amino acid (Sigma-Aldrich, U.S.A) and 1% penicillin-streptomycin (Sigma-Aldrich). Both cell lines were cultured at 37°C in a 5% CO_2_ humidified environment. DMEM/F-12 without penicillin-streptomycin was used for the co-culture assays. DMEM/F-12 without phenol red was used for the ROS detection assays.

### T84 and Caco-2 cells infection assays

Human IECs were counted using hemocytometer (Thermo Fisher Scientific, U.S.A) and for general infection assays, approximately 10^5^ cells were seeded in 24-well tissue culture plates 7 days prior to initiation of the *C. jejuni* infection. The plates were incubated at 37°C in a 5% CO_2_ atmosphere. For Western blotting, approximately 2 × 10^5^ cells were seeded in 6-well tissue culture plates. Prior to the infection, IECs were washed with phosphate-buffered saline (PBS; Thermo Fisher Scientific) three times and the medium was replaced with DMEM/F-12 without penicillin-streptomycin. *C. jejuni* strains grown on CBA plates for 24 hours were resuspended in PBS and bacterial suspension with appropriate OD_600_ were then incubated with IECs for various time periods giving a multiplicity of infection (MOI) of 200:1. In some experiments, T84 and Caco-2 cells were pre-treated with 10 μM DPI for 1 hour, washed three times with PBS and then infected with *C. jejuni*.

### DCFDA measurement of intracellular reactive oxygen species (ROS)

To analyse the levels of intracellular ROS production in human IECs under experiments conditions, DCFDA Cellular ROS Detection Assay Kit (Abcam U.K) was used according to the manufacturer’s instructions. Briefly, IECs grown in 96-well cell culture plates were washed three times with PBS and incubated with *C. jejuni* for 3 or 24 hours (MOI 200:1). For positive controls, IECs were treated with 500 μM H_2_O_2_ for 45 minutes. 45 minutes prior to completion of the infection, 100 μM 2’,7’-dichlorofluorescin diacetate (DCFDA) was added into each well giving a final concentration of 50 μM. After *C. jejuni* infection, the fluorescence was detected using SpectraMax M3 Multi-Mode Microplate Reader (Molecular Devices, U.S.A) with 485 nm excitation and 535 nm emission.

### Measurement of extracellular H_2_O_2_

Amplex^®^ Red Hydrogen Peroxide/Peroxidase Assay Kit (Invitrogen, U.S.A) was used to measure extracellular H_2_O_2_ in culture media after incubation with *C. jejuni*. Briefly, Amplex^®^ Red reagent (10-acetyl-3,7-dihydroxyphenoxazine) with horseradish peroxidase (HRP) reacts with H_2_O_2_ in a 1:1 stoichiometry producing a fluorescent product called resorufin. After incubation with *C. jejuni* for 3 or 24 hours, 100 μl of culture media was transferred to a 96-well plate and 100 μl of reaction mixture containing 50 μM Amplex^®^ Red reagent was added followed by incubation for 10 minutes at 37°C under microaerobic condition. Using SpectraMax M3 Multi-Mode Microplate Reader, fluorescence was measured at 530 nm excitation and 590 nm emission.

### Real time-quantitative polymerase chain reaction (qRT-PCR) analysis

For qRT-PCR, RNA was isolated from infected and uninfected IECs using PureLink™ RNA Mini Kit (Thermo Fisher Scientific) and contaminating DNA was removed using TURBO DNA-free kit (Ambion, U.S.A) according to manufacturer’s instructions. Concentration and purity of RNA samples were determined in a NanoDrop ND-1000 spectrophotometer (Thermo Fisher Scientific). 400 ng of RNA per sample were first denatured at 65°C for 5 minutes and snap cooled on ice. Complementary DNA (cDNA) was generated with random hexamers and SuperScript III Reverse Transcriptase (Thermo Fisher Scientific). Each reaction had 10 μl of SYBR Green PCR Master Mix (Applied Biosystems, U.S.A), 1 μl of primer (20 pmol), 1 μl of cDNA and 10 μl of HyClone™ water (Thermo Fisher Scientific). The sequence of each primer is described in Table S2. All reactions were run in triplicate on an ABI-PRISM 7500 instrument (Applied Biosystems) and expression levels of all target genes were normalised to *gapdh* expression determined in the same sample. Relative expression changes were calculated using the comparative threshold cycle (C_T_) method (Pfaffl, 2001). A minimum of three biological replicates were always analysed, each in technical triplicate.

### Semi-quantitative reverse transcription (RT-PCR) analysis

Each PCR reaction had 50 μl of FasTaq PCR master mix (Qiagen, Netherlands), 2 μl of primer (0.4 nmol) described in Table S2 and 1.5 μl of cDNA. For PCR reactions Tetrad-2 Peltier thermal cycler (Bio-Rad, U.K) was used. One cycle of PCR programme performs 95°C for 15 seconds after 2 minutes in the first cycle, annealing at 50°C for 20 seconds, and extension at 72°C for 30 seconds. Total 36 cycles were repeated. The PCR products were loaded on the 1% agarose gel and the gel was running for 1 hour at 120V. The gel was imaged using G:BOX Chime XRQ (Syngene, U.S.A). Quantification of relative mRNA level was performed using ImageJ software (Schneider, Rasband, & Eliceiri, 2012).

### SDS-PAGE and Western blot analysis

After infection, IECs were washed three times with PBS and lysed with cold RIPA lysis and extraction buffer (Thermo Fisher Scientific) with cOmplete™ Mini EDTA-free Protease Inhibitor Cocktail (Roche, Switzerland) and cleared by centrifugation (4 °C, 13,000 × g, 20 min). Protein concentration was determined using Pierce™ Bicinchoninic acid (BCA) Protein Assay Kit (Thermo Fisher Scientific) according to the manufacturer’s instructions. Afterward, samples were diluted to a desired concentration in HyClone™ water and 4X Laemmli sample buffer (Sigma-Aldrich) and incubated for 5 minutes at 95°C. Equal amounts of protein samples were separated using 4-12% NuPAGE™ Bis-Tris gel in 1× NuPAGE™ MES buffer or MOPS buffer (Thermo Fisher Scientific). Proteins were transferred from the gel using the iBlot® 2 transfer stacks (Life Technologies, U.S.A) using the iBlot® Gel Transfer Device (Invitrogen). These stacks were integrated with nitrocellulose transfer membrane. After the transfer, membranes were blocked with 1X PBS containing 2% (w/v) milk. Membranes were then probed with primary antibodies overnight as described previously (Elmi et al., 2016). The following primary antibodies were used; GAPDH (ab181602; Abcam); Nox1 (ab101027; Abcam) or Nox1 (NBP-31546; Novus Biologicals). Blots were developed using LI-COR infrared secondary antibody (IRDye 800CW Donkey anti-rabbit IgG) and imaged on a LI-COR Odyssey Classic (LI-COR Biosciences, U.S.A). Quantification of relative protein levels were performed using ImageJ software (Schneider et al., 2012).

### Detection of GTP-bound Active Rac1

The levels of active GTP-bound Rac1 were measured by using Rac1 G-LISA kit (Cytoskeleton Inc., U.S.A) according to the manufacturer’s instructions. Briefly, before the infection, IECs were incubated with reduced serum (0.1% FBS) for 24 hours. Infected or uninfected human IECs were washed with 1X PBS and lysed using the supplied 1X Lysis Buffer. Cell lysates were centrifuged for 1 minute at 10,000 x g at 4°C and adjusted to 1 mg/ml for the further process of the assay. As a positive control, constitutively active Rac1 (RCCA) was provided in the kit. Three biological replicates were conducted in all experiments, along with two technical replicates for each assay.

### Inhibition of Nox1 with diphenyleneiodonium chloride (DPI)

A stock solution of 3.25 mM DPI (Sigma-Aldrich) in dimethyl sulfoxide (DMSO; Sigma Aldrich) was prepared and stored at -20°C. For treatment, the DPI stock solution was diluted to 10 μM DPI in culture media without antibiotics, then incubated with IECs for 1 hour at 37°C in a 5% CO_2_ atmosphere. After treatment, IECs were washed with PBS for three times before co-incubation with *C. jejuni* for various time points.

### Small interfering (si) RNA transfection

On the day of reverse transfection, 500 μl of Caco-2 cells (10^5^ cells/ml) were seeded in 24-well plates and treated for 24 hours with 30 pmol siRNA from either Nox1 siRNA (sc-43939; Santa Cruz Biotechnology, Inc, U.S.A) or Ambion® Silencer Negative Control #1 siRNA (Invitrogen) for the negative control. For preparation of siRNA transfection reagent complex, 3 μl of 10 μM stock siRNA was diluted with 100 μl of Opti-MEM^®^ Reduced-Serum Medium (Thermo Fisher Scientific) and mixed with 1.5 μl of Lipofectamine^®^ RNAiMAX Transfection Reagent (Thermo Fisher Scientific). After 24 hours transfection, media was replaced with DMEM/F-12 containing 10% FBS. After additional 48 or 72 hours, RNA and protein were extracted to check efficacy of transfection.

### Adhesion, invasion and intracellular survival assay

Adherence, invasion and intracellular assays were performed as described previously with minor modifications (Gundogdu et al., 2011). T84 and Caco-2 cells seeded in a 24-well plate were washed three times with PBS and treated with 10 μM DPI for 1 hour or transfected with Nox1 siRNA as described in above. Then IECs were inoculated with *C. jejuni* with OD_600_ 0.2 at a MOI of 200:1 and incubated for 3 hours at 37°C in 5% CO_2_. For the interaction (adhesion and invasion) assay, monolayers were washed three times with PBS to remove unbound extracellular bacteria and then lysed with PBS containing 0.1% (v/v) Triton X-100 (Sigma-Aldrich) for 20 min at room temperature. The cell lysates were diluted and plated on blood agar plates to determine the number of interacting bacteria (CFU/ml).

Invasion assays were performed by additional step of treatment of gentamicin (150 μg/ml) for 2 hours to kill extracellular bacteria, washed three times with PBS, lysed and plated as described above. For intracellular survival assays, after infection with *C. jejuni* for 3 hours, T84 and Caco-2 cells were treated with gentamicin (150 μg/ml) for 2 hours to kill extracellular bacteria followed by further 18 hours incubation with gentamicin (10 μg/ml). Cell lysis and inoculation were performed as described above.

### Cytotoxicity assay with trypan blue exclusion methods

After treatment with DPI and gentamicin or transfection with siRNA as previously described, IECs were washed three times with PBS and were detached using trypsin-EDTA (Thermo Fisher Scientific) and resuspended with culture media. 50 μl of cell suspension were added into 50 μl of 0.4% Trypan Blue solution (Thermo Fisher Scientific) and the numbers of viable and dead cells were counted using hemocytometer under microscope.

### *Campylobacter jejuni* viability test with DPI treatment

T84 cells were treated with 10 μM DPI for 1 hour and the cells were washed three times with PBS. After DPI treatment for 1 hour, *C. jejuni* strains (OD_600_ 0.2) were co-incubated for 1 hour with PBS from the last wash. After incubation, serial dilution was performed and each dilution was spotted on to blood agar plates. The plates were incubated under microaerobic condition at 37°C for 48 hours. CFU of each spot was recorded.

### Statistical analysis and graphing

At least three biological replicates were performed in all experiments. Each biological replicate was performed in three technical replicates. For statistical analysis and graphing, GraphPad Prism 8 for Windows (GraphPad Software, U.S.A) was used. One sample *t*-test or unpaired *t*-test were used to compare two data sets for significance with * indicating *p* < 0.05, ** indicating *p* < 0.01, *** indicating *p* < 0.001, and **** indicating *p* < 0.0001.

## Results

### *Campylobacter jejuni* modulates intracellular and extracellular ROS in T84 and Caco-2 cells in a time- and strain-dependent manner

As *C. jejuni* possesses distinct physiological characteristics compared to more studied enteric pathogens, we assessed the ability of three distinct *C. jejuni* strains to modulate intracellular and extracellular ROS in T84 and Caco-2 cells (Burnham & Hendrixson, 2018). We observed strain-specific ROS modulation at 3- and 24-hours post-infection (Figure 1). All three *C. jejuni* strains reduced the levels of intracellular ROS in T84 and Caco-2 cells compared to the uninfected control (Figure 1A, 1B, 1E, 1F). A similar pattern was observed for extracellular ROS where all *C. jejuni* strains reduced the levels of extracellular ROS in T84 and Caco-2 cells compared to the uninfected control (Figure 1C, 1D, 1G, 1H). A distinct pattern was observed when assessing levels of extracellular ROS for *C. jejuni* 81-176 strain at 3 hours post-infection (Figure 1C and 1G). At this early time point, extracellular ROS is increased in T84 and Caco-2 cells infected with *C. jejuni* 81-176, although we observed similar reduced levels of ROS at 24 hours. These results indicate a strain and time-specific pattern linking the ability of different *C. jejuni* strains to modulate intracellular and extracellular ROS levels in T84 and Caco-2 cells.

**FIGURE 1.**
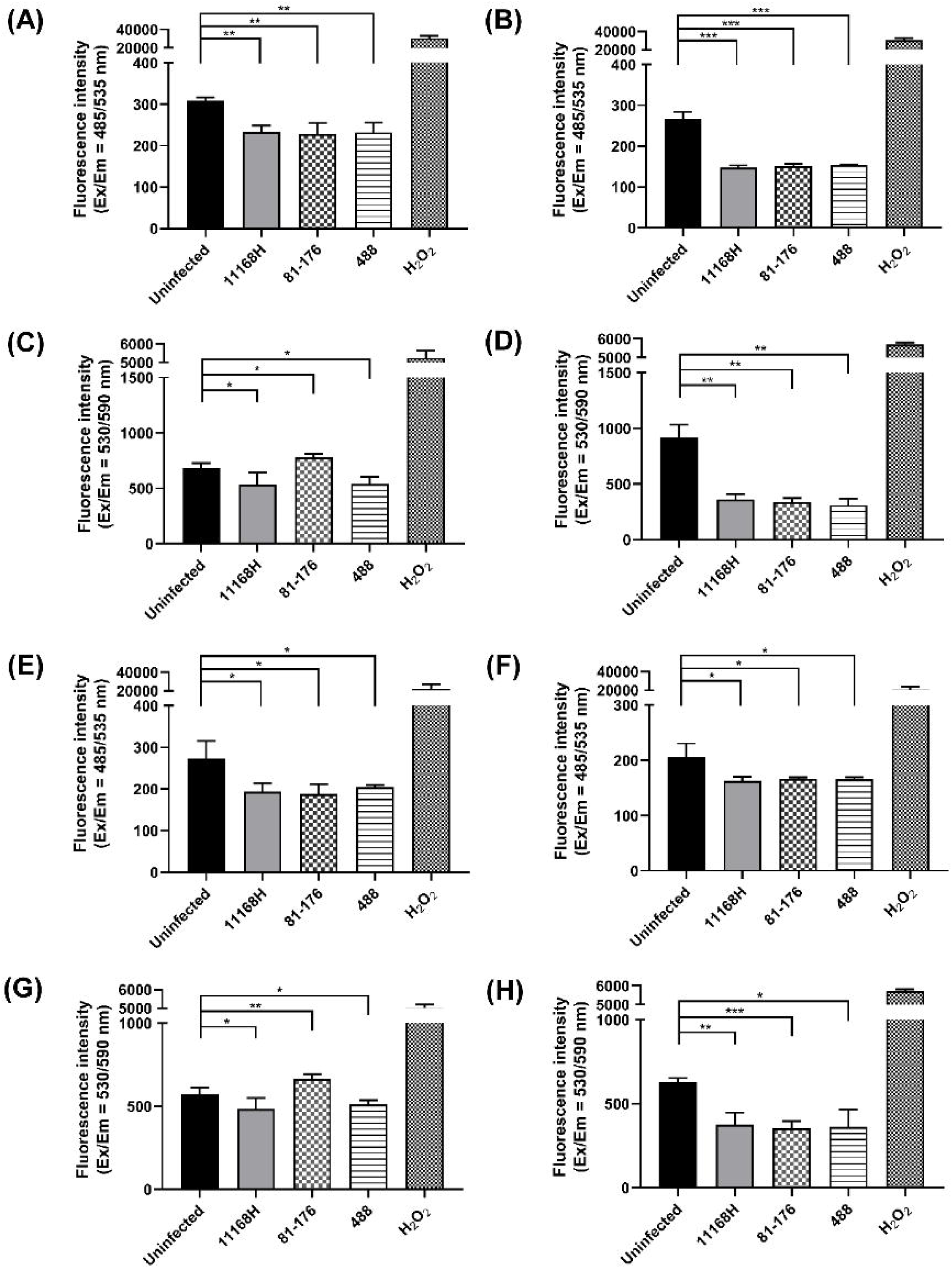
Detection of intracellular and extracellular ROS in T84 and Caco-2 cells after infection with *C. jejuni* 11168H, 81-176 or 488 strains. Intracellular ROS in T84 cells after infection with *C. jejuni* for (A) 3 hours or (B) 24 hours and extracellular ROS from T84 cells after infection with *C. jejuni* for (C) 3 hours or (D) 24 hours were measured. Intracellular ROS in Caco-2 cells after infection of *C. jejuni* for (E) 3 hours or (F) 24 hours and extracellular ROS from Caco-2 cells after infection for (G) 3 hours or (H) 24 hours were measured. For detection of intracellular ROS, DCFDA was used. For detection of extracellular ROS, Amplex^®^ Red reagent with HRP were used. H_2_O_2_ was used as a positive control. Experiments were repeated in three biological and three technical replicates. Asterisks denote a statistically significant difference (* = *p* < 0.05; ** = *p* < 0.01; *** = *p* < 0.001).

### *Campylobacter jejuni* modulates intracellular and extracellular ROS in T84 and Caco-2 cells via the downregulation of Nox1 complex

Given the observed modulation of intracellular and extracellular ROS in T84 and Caco-2 cells, we next explored the mechanism by which *C. jejuni* strains orchestrate ROS modulation. We analysed the transcription and translation of Nox1 which is the main source of ROS production in IECs (Brandes et al., 2014; Sumimoto et al., 2005). As shown in Figure 2, Nox1 transcription and translation levels were significantly reduced in both T84 (Figure 2A) and Caco-2 cells (Figure 2B) infected with *C. jejuni* when compared to uninfected cells. Notably, at 24 hours post-infection, mRNA levels of Nox1 in T84 cells are significantly reduced compared with *C. jejuni*-infected Caco-2 cells. We measured the relative levels of mRNA between T84 and Caco-2 cells and identified T84 cells expressed a higher basal level of Nox1 mRNA compared to Caco-2 cells (Figure 2C). As a result of this higher basal level of Nox1 mRNA in T84 cells we validated our qRT-PCR data using RT-PCR where less expression of Nox1 in *C. jejuni*-infected T84 cells was observed (Figure 2D and 2E). Reduction in the translational level of Nox1 in *C. jejuni*-infected T84 cells was confirmed independently by Western blotting (Figure 2F and 2G).

**FIGURE 2.**
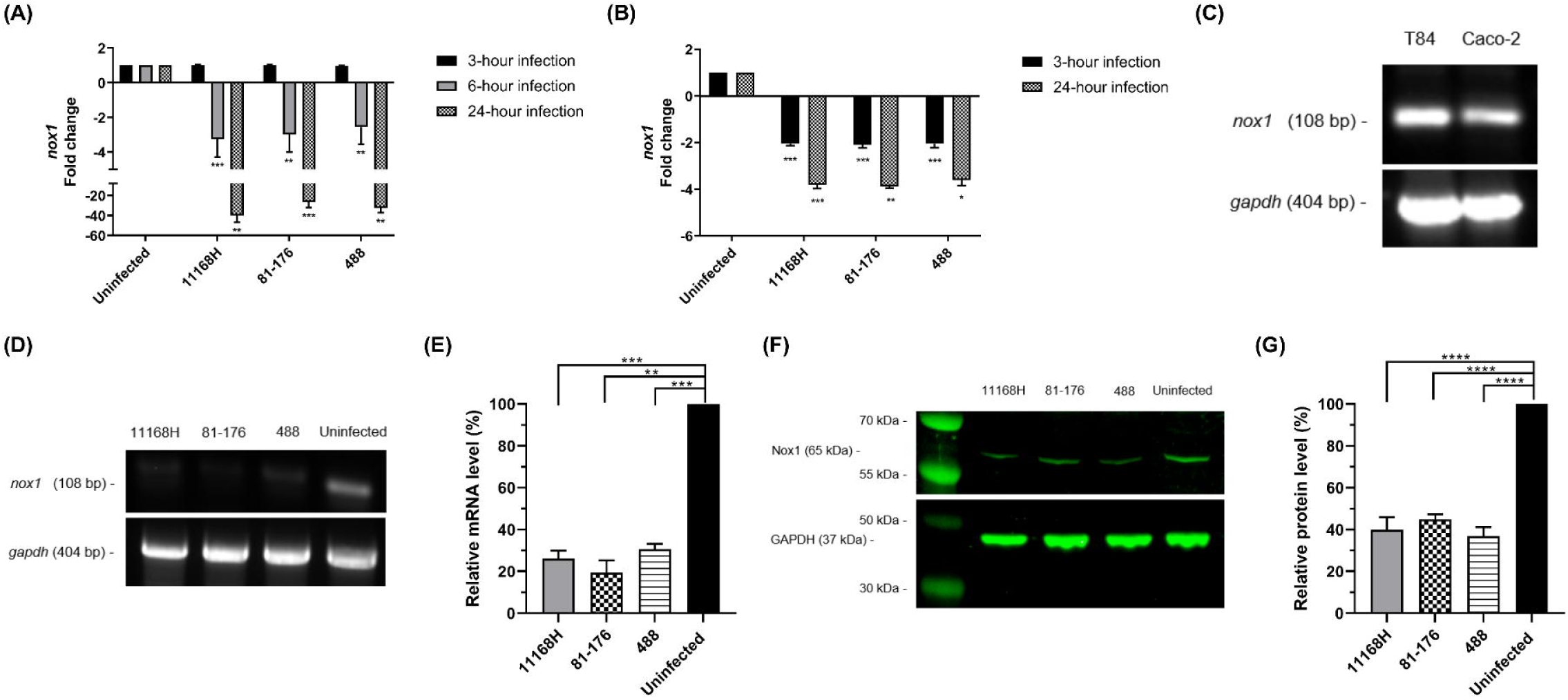
*C. jejuni* modulates Nox1 expression in T84 and Caco-2 cells. qRT-PCR showing expression of Nox1 in (A) T84 and (B) Caco-2 cells. (C) RT-PCR showing expression of Nox1 in uninfected T84 and Caco-2 cells. *gapdh* was used as an internal control. (D) RT-PCR showing expression of Nox1 in T84 cells infected with *C. jejuni* for 24 hours and (E) relative mRNA levels as a percentage from RT-PCR data. (F) Western blotting showing Nox1 in T84 cells infected with *C. jejuni* for 24 hours and (G) relative protein level as a percentage from Western blotting. Asterisks denote a statistically significant difference (* = *p* < 0.05; ** = *p* < 0.01; *** = *p* < 0.001).

### *Campylobacter jejuni* modulates activity of small GTPase Rac1 in T84 and Caco-2 cells in a time-dependent manner

To gain further insight into the mechanism that leads to *C. jejuni* modulation of ROS in T84 and Caco-2 cells, we examined the ability of *C. jejuni* to activate Rac1, a member of the Rho family of small GTPases. Although Rac1 is implicated in Nox1 activation in several eukaryotic cell lines (Nisimoto et al., 2008; Ueyama et al., 2006), the contribution of Rac1 in *C. jejuni*-mediated Nox1 modulation is unknown. As shown in Figure 3, Rac1 is an integral part of the Nox1 complex. Given that *C. jejuni* activates Rac1 in human INT 407 cells via *Campylobacter* invasion antigen D (CiaD) (Krause-Gruszczynska et al., 2007; Negretti et al., 2021), and that Rac1 supports Nox1 activity only in its GTP-bound active form, we examined if downregulation of Nox1 is linked to GTPase Rac1 by *C. jejuni*. Interestingly, *C. jejuni* 11168H strain induced Rac1 1- and 3-hours after infection in T84 cells (Figure 4A). After 24 hours infection, Rac1 activity was reduced (though not statistically significant; *p* = 0.0714) (Figure 4A). Similarly, *C. jejuni* 11168H induced Rac1 activity after 1 hour infection in Caco-2 cells (Figure 4B). However, this activity was reduced after 3- and 24-hours infection (Figure 4B). These results suggest that the downregulation of Nox1 by *C. jejuni* is inversely correlated with an increase in Rac1 GTPase activity.

**FIGURE 3.**
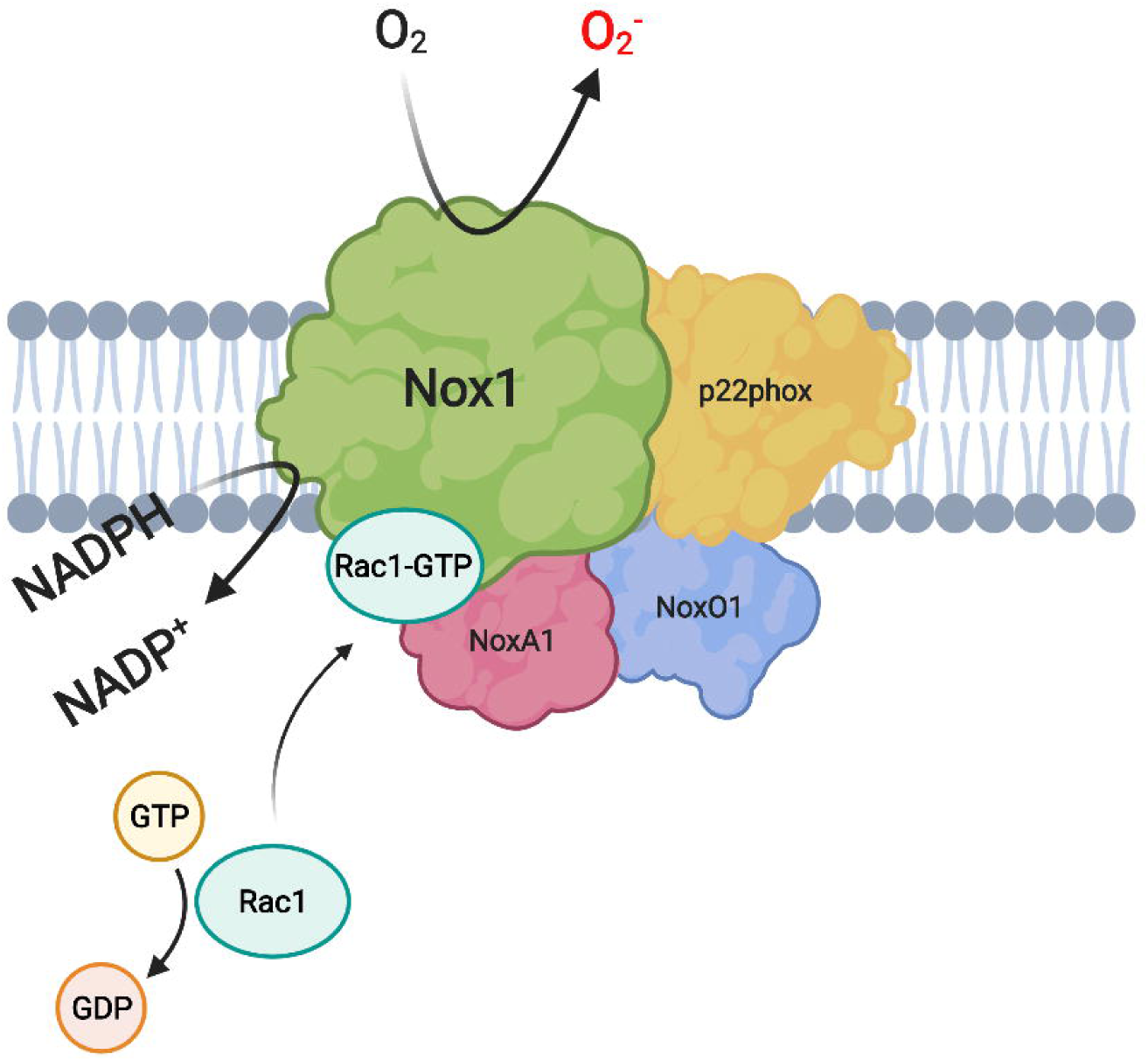
Proposed structure of the Nox1 complex consisting of Nox1, p22phox, GTP-bound Rac1, NoxA1 and NoxO1. p22phox and other subcellular subunits are assembled to activate catalytic subunit Nox1 which results in the generation of O_2_^−^ by oxidising NADPH (Brandes et al., 2014). Created with BioRender.com

**FIGURE 4.**
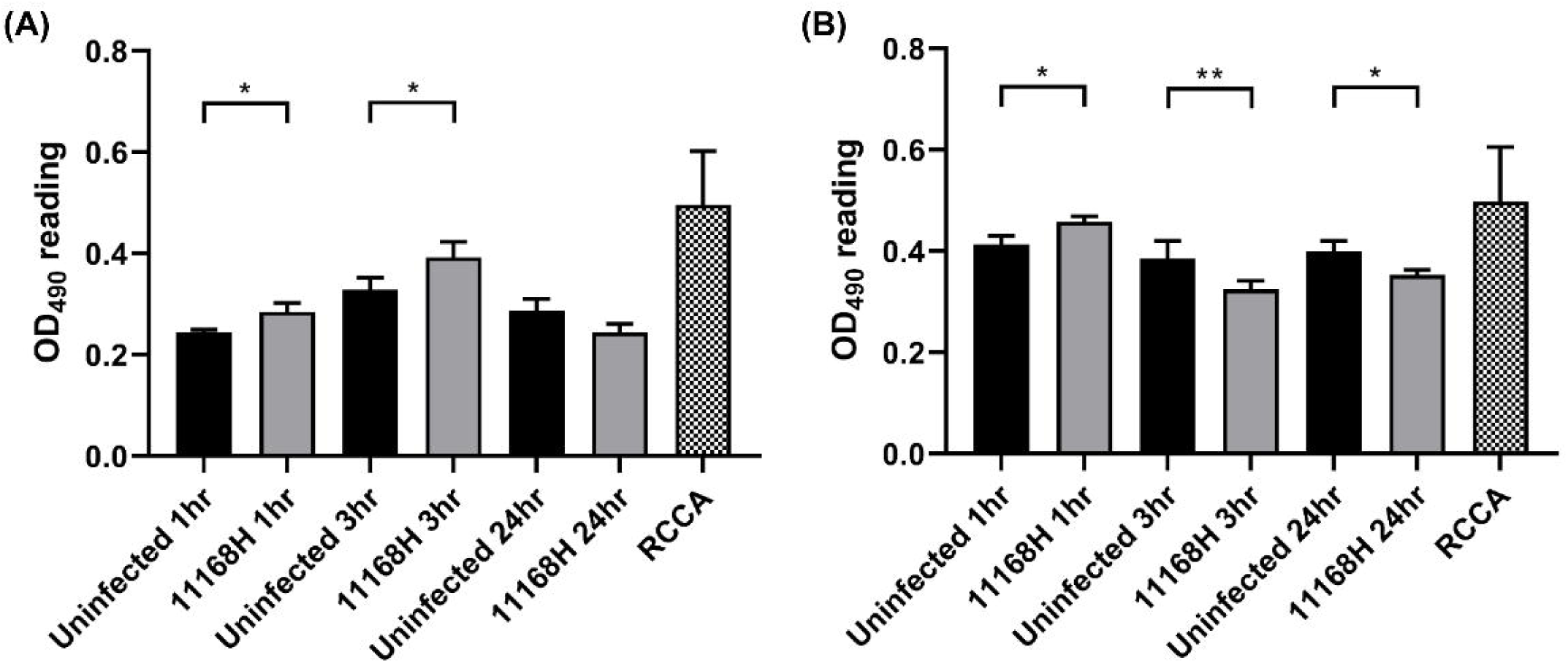
*C. jejuni* modulates activity of small GTPase Rac1 in T84 and Caco-2 cells. (A) T84 and (B) Caco-2 cells were infected with *C. jejuni* 11168H strain for 1, 3, and 24 hours and the activation of small GTPase Rac1 in each time point was measured. Constitutively active Rac1 (RCCA) was used as a positive control. Experiments were repeated in three biological and three technical replicates. Asterisks denote a statistically significant difference (* = *p* < 0.05, ** = *p* < 0.001).

### *Campylobacter jejuni* modulates transcription of antioxidant-related genes in T84 and Caco-2 cells

To gain further insight into the ability of *C. jejuni* to modulate intracellular and extracellular ROS in T84 and Caco-2 cells, we sought to understand if *C. jejuni* modulates the expression of two important antioxidant genes, superoxide dismutase 1 (*sod1*) and catalase (*cat*). Sod1 decomposes O_2_^-^ to H_2_O_2_, and Cat breaks down H_2_O_2_ to H_2_O and O_2_ (Aviello & Knaus, 2017). Intriguingly, as shown in Figure 5A and 5B, there is a significant downregulation of the mRNA levels of *cat* and *sod1* at 24 hours post-infection in T84 cells. A similar pattern was observed when compared with Caco-2 cells where the expression of *cat* and *sod1* at 24 hours post-infection is significantly downregulated (Figure 5C and 5D). In contrast, the expression of *cat* and *sod1* at 3 hours post-infection is unaffected. These results may indicate *C. jejuni*-mediated reduction in intracellular and extracellular ROS is independent of modulation of *cat* and *sod1*.

**FIGURE 5.**
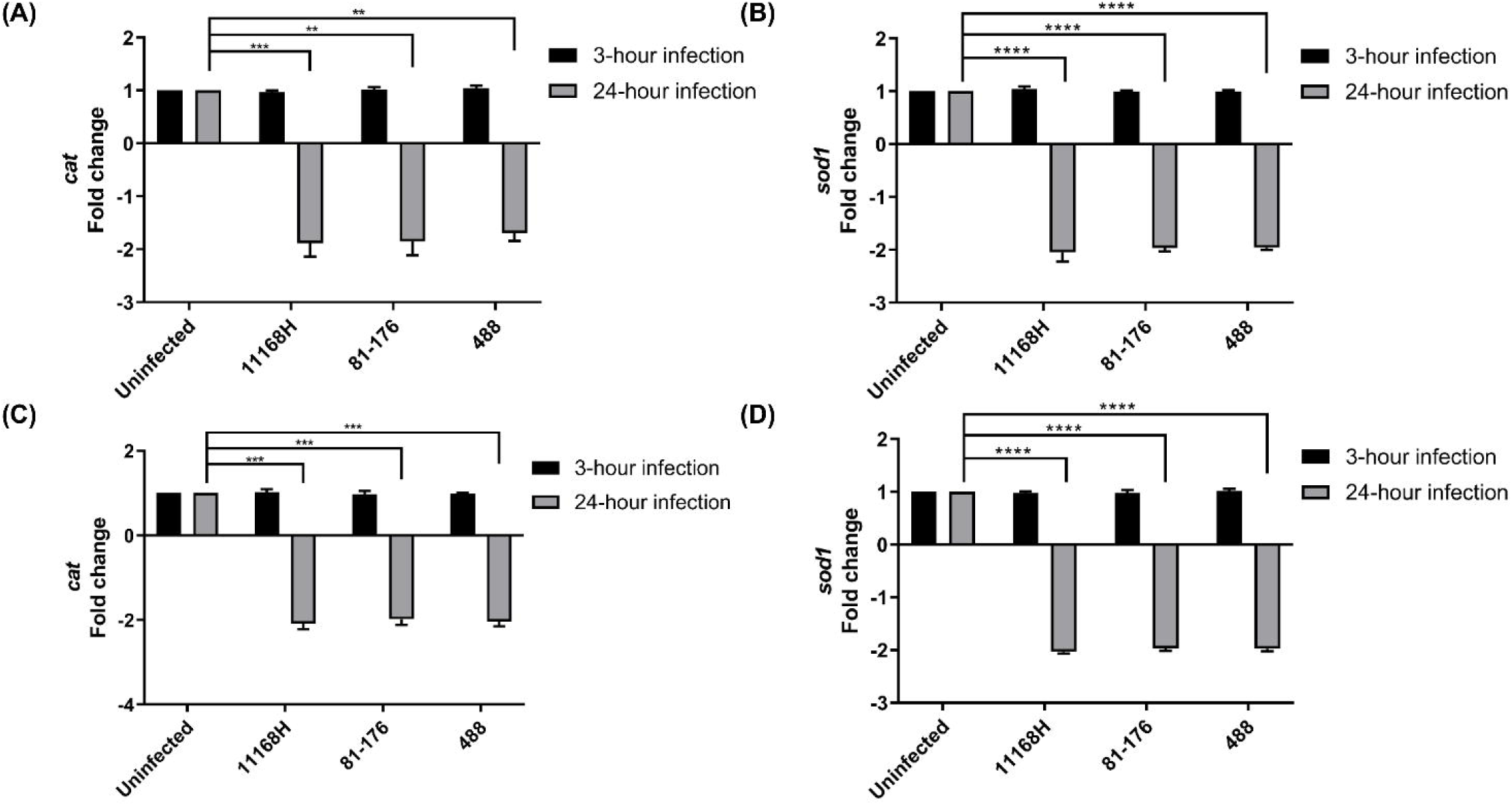
qRT-PCR showing expression of human catalase (*cat*) and superoxide dismutase 1 (*sod1*) in T84 and Caco-2 cells. (A, B) T84 and (C, D) Caco-2 cells were infected with *C. jejuni* for 3- or 24-hours and transcriptional levels of *cat* and *sod1* were measured. *gapdh* was used as an internal control. Experiments were repeated in three biological and three technical replicates. Asterisks denote a statistically significant difference (** = *p* < 0.01; *** = *p* < 0.001; **** = *p* < 0.0001).

### Chemical inhibition of Nox1 activity by DPI impairs *Campylobacter jejuni* interaction, invasion and intracellular survival of T84 and Caco-2 cells *in vitro*

Having established that *C. jejuni* significantly reduced the transcription and translation of Nox1 in T84 and Caco-2 cells in a time-dependent manner, and that Rac1 is not only known as a key component of the Nox1 complex, but also implicated in cell dynamic morphology (Nisimoto et al., 2008; Ueyama et al., 2006), we hypothesised Rac1-mediated Nox1 might modulate membrane ruffling and cytoskeleton rearrangement which might in turn affect *C. jejuni* interaction with IECs. Therefore, we investigated the role of Nox1 in *C. jejuni* interaction, invasion and intracellular survival in IECs by transiently pre-treating T84 and Caco-2 cells with DPI (10 μM) which is known to inhibit activity of flavoenzymes including Nox complex (Riganti et al., 2004). First, we demonstrated that DPI reduced extracellular ROS in T84 and Caco-2 cells (Figure S1). As shown in Figure 6A, 6C, 6E, pre-treatment of T84 cells by DPI significantly reduced the ability of *C. jejuni* to interact, invade, and survive intracellularly in T84 cells. Similarly, as shown in Figure 6B, 6D and 6F, *C. jejuni* infected with DPI-treated Caco-2 cells showed significant reduction in interaction, invasion, and intracellular survival compared to untreated Caco-2 cells. Since our data revealed *C. jejuni* reduced interaction, invasion and intracellular survival between the control and DPI-treated T84 and Caco-2 cells, we next evaluated the viability of *C. jejuni*, T84 and Caco-2 cells co-incubated with DPI. Treatment with DPI did not affect viability of IECs (Figure S2) or *C. jejuni* (Figure S3). Thus, our observations suggest further inhibition of Nox1 with DPI is detrimental to *C. jejuni* interaction, invasion and intracellular survival in IECs.

**FIGURE 6.**
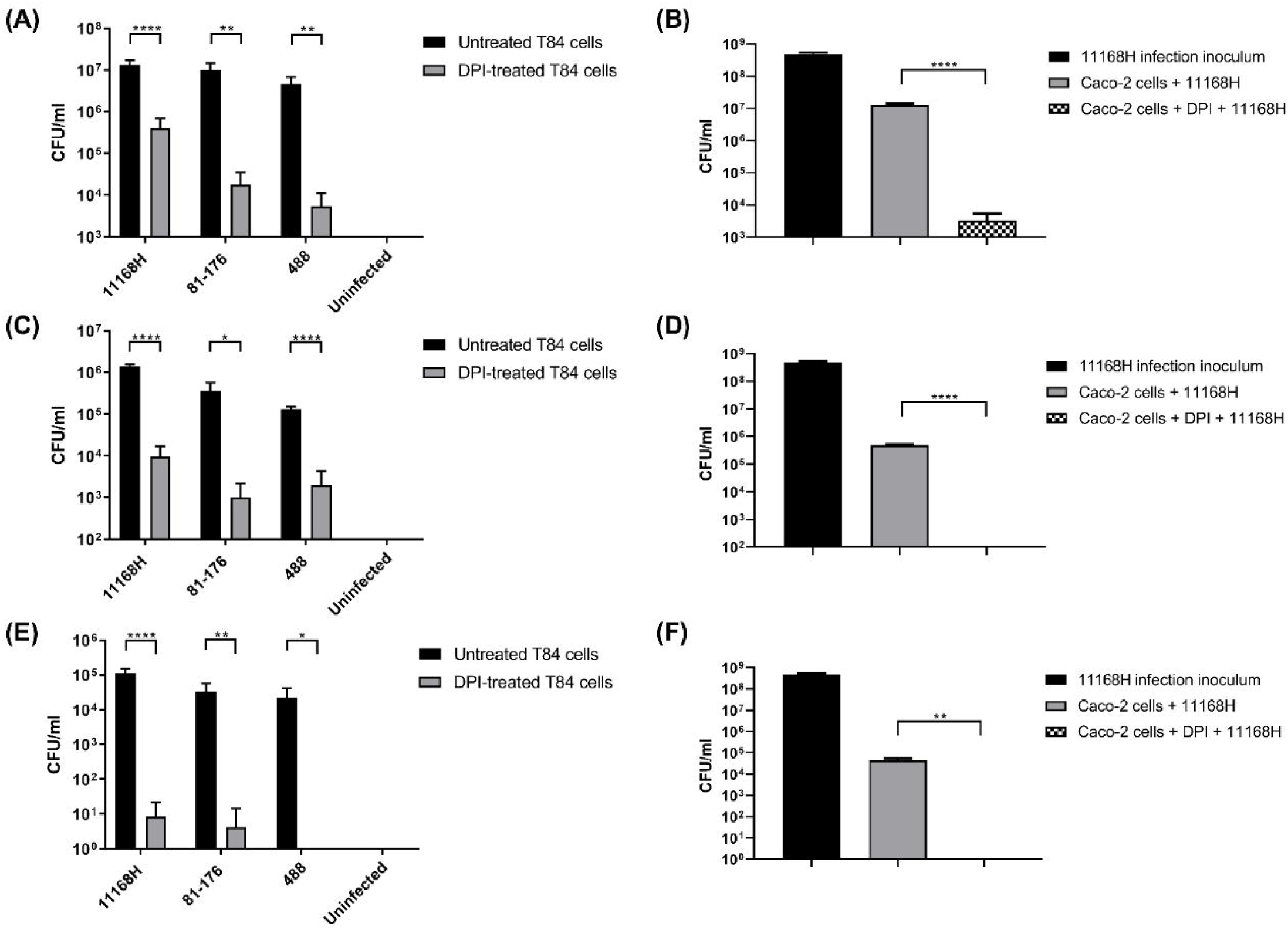
The effect of DPI on *C. jejuni* interaction, invasion and intracellular survival. T84 and Caco-2 cells were pre-treated with 10 μM of DPI for 1 hour and infected with C. jejuni for 3 hours. (A) T84 and (B) Caco-2 cells were washed with PBS and lysed, and the numbers of interacting bacteria were assessed. (C, D) For invasion assay, after infection with C. jejuni, *IECs* were incubated with gentamicin (150 μg/ml) for 2 hours to kill extracellular bacteria and then lysed, and the numbers of intracellular bacteria were assessed. (E, F) For intracellular survival assay, 2 hours gentamicin treatment was followed by further incubation with gentamicin (10 μg/ml) for 18 hours. Then cells were lysed, and the number of intracellular bacteria were assessed. Experiments were repeated in three biological and three technical replicates. Asterisks denote a statistically significant difference (* = p < 0.05; ** = p < 0.01; **** = p < 0.0001).

### Nox1 silencing by siRNA impairs *Campylobacter jejuni* interaction, invasion and intracellular survival in Caco-2 cells *in vitro*

As DPI is a pan-Nox inhibitor, we silenced Nox1 expression in Caco-2 cells by delivering specific small interfering RNA (siRNA) into cultured Caco-2 cells. We used siRNA sequence which target regions of Nox1 for silencing. As a negative control, we used a non-targeting scrambled RNA sequence which is not complementary to the Nox1 mRNA. As shown in Figure 7A and 7B, transcriptional and translational levels of Nox1 were significantly decreased in cells treated with Nox1 siRNA, relative to that in mock-treated Caco-2 controls. We further confirmed reduced activity of Nox1 by demonstrating significant reduction in extracellular ROS (Figure 7C). We showed that Nox1 siRNA transfection did not affect viability of Caco-2 cells (Figure S4). Based on these results, we further investigated interaction, invasion and intracellular survival of *C. jejuni* within Caco-2 cells (Figure 7D, 7E and 7F). Our result showed significant decrease in *C. jejuni* interaction, invasion and intracellular survival when compared to non-transfected controls. This result highlights a correlation between reduced Nox1 expression with a reduction in *C. jejuni* infection. Taken together, our results demonstrate that Nox1 is a critical host factor for *C. jejuni* interaction, invasion, and intracellular survival.

**FIGURE 7.**
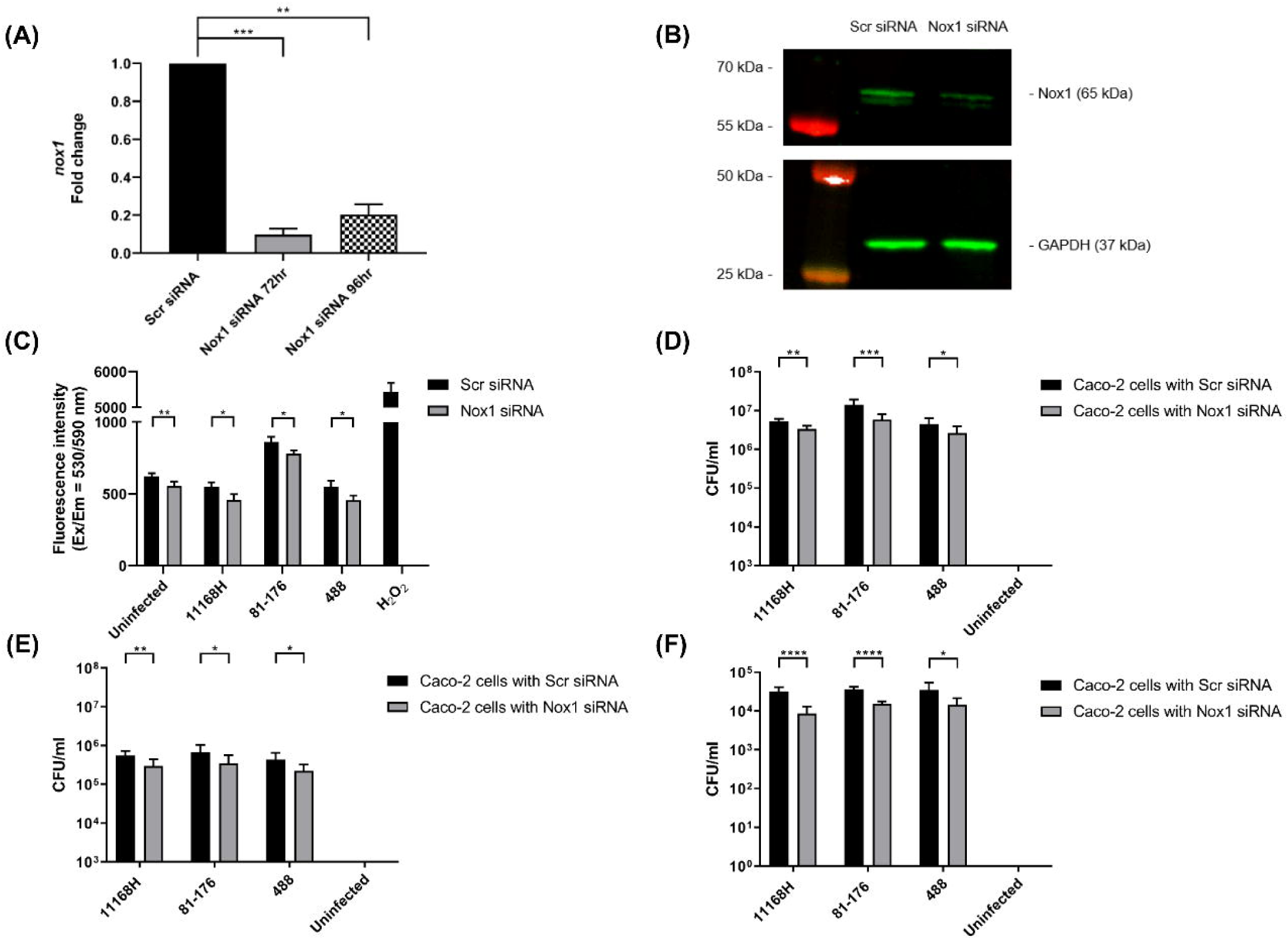
The effect of Nox1 silencing on *C. jejuni* interaction, invasion and intracellular survival. Caco-2 cells were transfected with Nox1 siRNA or scrambled siRNA (Scr siRNA). (A) qRT-PCR showing expression of Nox1 after siRNA transfection. (B) Western blotting showing expression of Nox1 after 72 hours siRNA transfection. (C) Detection of extracellular ROS from Caco-2 cells after 72 hours siRNA transfection followed by co-incubation of *C. jejuni* for 3 hours. (D) After 72 hours siRNA transfection followed by *C. jejuni* infection for 3 hours, Caco-2 cells were washed with PBS and lysed and the numbers of interacting bacteria were assessed or (E) for invasion assay, the cells were incubated with gentamicin (150 μg/ml) for 2 hours to kill extracellular bacteria and then lysed, and the numbers of intracellular bacteria were assessed. (F) For intracellular survival assay, 2 hours gentamicin treatment was followed by further incubation with gentamicin (10 μg/ml) for 18 hours. Then the cells were lysed, and the number of intracellular bacteria determined. Experiments were repeated in three biological and three technical replicates. Asterisks denote a statistically significant difference (* = *p* < 0.05; ** = *p* < 0.01; *** = *p* < 0.001; **** = *p* < 0.0001).

## Discussion

Upon infection, host cells induce a range of cellular responses to remove offending pathogens. However, bacterial pathogens often target host organelle(s), signalling pathway(s) or immune responses to evade host defence mechanisms (Escoll et al., 2016). Disruption of ROS production in host cells by bacterial pathogens has been previously reported (Gallois, Klein, Allen, Jones, & Nauseef, 2001; Vareechon, Zmina, Karmakar, Pearlman, & Rietsch, 2017). *S*. Typhimurium pathogenicity island-2 encoding Type III Secretion System (T3SS) inhibits ROS production in human macrophages by preventing Nox2 assembly (Antoniou et al., 2018; Gallois et al., 2001). In addition, *Pseudomonas aeruginosa* T3SS effector, ExoS disrupts ROS production in human neutrophils by ADP-ribosylating Ras and inhibiting its activity which is essential for Nox2 assembly (Vareechon et al., 2017).

We have characterised the ability of distinct *C. jejuni* strains to modulate intracellular and extracellular ROS from human IECs *in vitro*. ROS production by human IECs is a major defence mechanism, yet how *C. jejuni* evades ROS remains unclear. Our work establishes that in contrast to other enteric pathogens, *C. jejuni* uses a different mechanism involving downregulation of Nox1 expression to modulate ROS in human IECs (den Hartog et al., 2016; Elatrech et al., 2015; Kawahara et al., 2005; Kawahara et al., 2016). We examined three different *C. jejuni* strains using two different human IECs and showed that *C. jejuni* strains modulate intracellular and extracellular ROS from human IECs via the differential regulation of the transcription and translation of Nox1 which is a major ROS source in IECs (Aviello & Knaus, 2017). Interestingly, a previous study demonstrated that *C. jejuni* 81-176 induces extracellular ROS production through Nox1 activation in human ileocecal adenocarcinoma derived HCT-8 cells (Corcionivoschi et al., 2012). To further understand the implications of *C. jejuni* transcriptional and translational downregulation of Nox1 in T84 and Caco-2 cells, we revealed similarities with some other enteropathogens, and also differences amongst others including the *C. jejuni* strain 81-176 (den Hartog et al., 2016; Elatrech et al., 2015; Kawahara et al., 2005; Kawahara et al., 2016). Enteropathogens such as *E. coli, Salmonella* spp., and *H. pylori* upregulate expression of Nox1 and ROS production in infected IECs (den Hartog et al., 2016; Elatrech et al., 2015; Kawahara et al., 2005; Kawahara et al., 2016). Our findings confirmed downregulation of ROS production by *C. jejuni* is strain dependent. In contrast to *C. jejuni* 11168H and 488 strains, *C. jejuni* 81-176 induced extracellular ROS in T84 and Caco-2 cells at 3 hours post-infection. Induction of extracellular ROS by *C. jejuni* 81-176 at this earlier infection time point was also observed previously (Corcionivoschi et al., 2012). We hypothesise *C. jejuni* 81-176 might have additional bacterial determinants which may induce host extracellular ROS independent of Nox1 modulation (*e*.*g*. the pVir and pTet plasmids which encode putative Type IV Secretion Systems (T4SS)) (Bacon et al., 2002; Batchelor, Pearson, Friis, Guerry, & Wells, 2004). We also noted a difference between the ability of *C. jejuni* strains to regulate expression of Nox1 in T84 and Caco-2 cells. This difference could be due to variations between the two cell lines. Caco-2 cells possess characteristic enterocytes whereas T84 cells possess characteristic colonocytes throughout differentiation (Devriese et al., 2017). In addition, previous studies have shown that reduced Nox1 mRNA was present in the ileum than in the colon of healthy patients suggesting there is a gradient in Nox1 expression from small intestine to large intestine (Schwerd et al., 2018). In our study, the lower expression of Nox1 mRNA detected in Caco-2 cells compared to T84 cells was also observed.

As ROS homeostasis in the GI tract is regulated by multiple antioxidant enzymes (Aviello & Knaus, 2017), *C. jejuni*-mediated modulation of Cat and Sod1 at the transcriptional level was investigated. Our data demonstrated C. *jejuni* strains did not affect transcriptional levels of *cat* and *sod1* in T84 and Caco-2 cells after 3 hours infection, but they significantly downregulated expression of both genes after 24 hours. To our knowledge, this is the first data on *C. jejuni* modulation of antioxidant-related genes in human IECs *in vitro*. Our observations imply *C. jejuni* might modulate intracellular or extracellular ROS after 3 hours infection without modulating expression of *cat* and *sod1*. These results also suggest that there could be additional mechanisms of *C. jejuni*-mediated reduction of ROS because *C. jejuni* was able to reduce ROS after 24 hours infection even though transcription levels of antioxidant-related genes *cat* and *sod1* were downregulated. However, we cannot disregard the possibilities that *C. jejuni* might secrete its own antioxidant-related proteins that may mitigate host cellular ROS and/or *C. jejuni* might induce expression of other host antioxidant genes such as mitochondrial superoxide dismutase (Sod2), extracellular superoxide dismutase (Sod3) and glutathione peroxidase (Aviello & Knaus, 2017).

Upon adhering to host cells, *C. jejuni* modulates small GTPase Rac1 resulting in actin filament reorganisation to promote invasion. Activation of Rac1 in human embryonic INT 407 cells was observed between 45 minutes and 4 hours after *C. jejuni* infection (Krause-Gruszczynska et al., 2007; Negretti et al., 2021). In accordance with previous studies, we demonstrated *C. jejuni* activates Rac1 at early infection time points. In contrast, a decrease of active Rac1 was detected at the later infection time point. Given the association of the active GTP-bound Rac1 and Nox1 activity, the early activation of Rac1 in IECs suggest that *C. jejuni* uses an intriguing system which we hypothesise could have temporally nonoverlapping mechanisms. The GTP-bound Rac1 observed in early time points may be linked to the requirement for *C. jejuni* to establish adhesion/invasion utilising a distinct mechanism in its infection cycle. Although the inactive GDP-bound Rac1 observed at the later time point of 24 hours, suggests *C. jejuni* clearly possesses yet to be discovered mechanisms that enable differential regulation of Nox1 relative to modulation of Rac1. We also observe the pattern of active GTP-bound Rac1 in Caco-2 cells that is different to T84 cells. Such a difference may be due to the signalling cues between the cells as well as *C. jejuni* preference to efficiently interact with individual cells by binding, invading, and intracellularly surviving from distinct states during its infection.

The impact of differential regulation of Nox1 on *C. jejuni* interaction, invasion and intracellular survival in human IECs remains unclear. Surprisingly, chemical inhibition of Nox1 significantly reduced the ability of *C. jejuni* to interact, invade, and survive intracellularly in T84 and Caco-2 cells. It is possible that DPI may inadvertently affect local cellular receptors that *C. jejuni* uses to bind human IECs. Since DPI is not a specific inhibitor of Nox1 (Riganti et al., 2004), we repeated these experiments using siRNA silencing of Nox1 which demonstrated similar findings, suggesting that Nox1 is indirectly necessary for *C. jejuni* interaction, invasion, and intracellular survival. Previous studies have demonstrated that DPI treatment reduced fibronectin expression in rat renal tubular epithelial cells (Rhyu et al., 2005), and a pan-Nox inhibitor APX-115 reduced fibronectin production in mesangial cells (Cha et al., 2017). As fibronectin has been demonstrated as a key host receptor that *C. jejuni* uses to bind and invade human IECs (Michael E. Konkel, Talukdar, Negretti, & Klappenbach, 2020), we hypothesise that silencing Nox1 might also affect expression of a key receptor fibronectin as is the case following DPI treatment, and this might be responsible for the reduced interaction and invasion of *C. jejuni* strains. However, the broader non-specificity of DPI and siRNA silencing experiments mean that there could be alternative mechanisms in play.

We have demonstrated that *C. jejuni* modulates intracellular and extracellular ROS in human T84 and Caco-2 cells. Our observations link *C. jejuni* ROS modulation to the transcriptional and translational downregulation of Nox1. These findings also point to a further role of Rac1 in Nox1 modulation and downstream interaction. Based on chemical inhibition and silencing of Nox1 expression and translation, our findings suggest an indirect role of Nox1 for adhesion, invasion and intracellular survival of *C. jejuni*. In this context, further understanding *C. jejuni* determinants that lead to ROS and/or Nox1 modulation in IECs will provide greater insights into how *C. jejuni* manipulate host defence mechanisms and cause diarrhoeal disease.

## ACKNOWLEDGEMENTS

We would like to acknowledge Marta Mauri for kind advice on siRNA transfection.

