## Supplementary Figure S1 for "*Campylobacter jejuni* modulates reactive oxygen species production and NADPH oxidase 1 expression in human intestinal epithelial cells"

**Supplementary Figure 1**

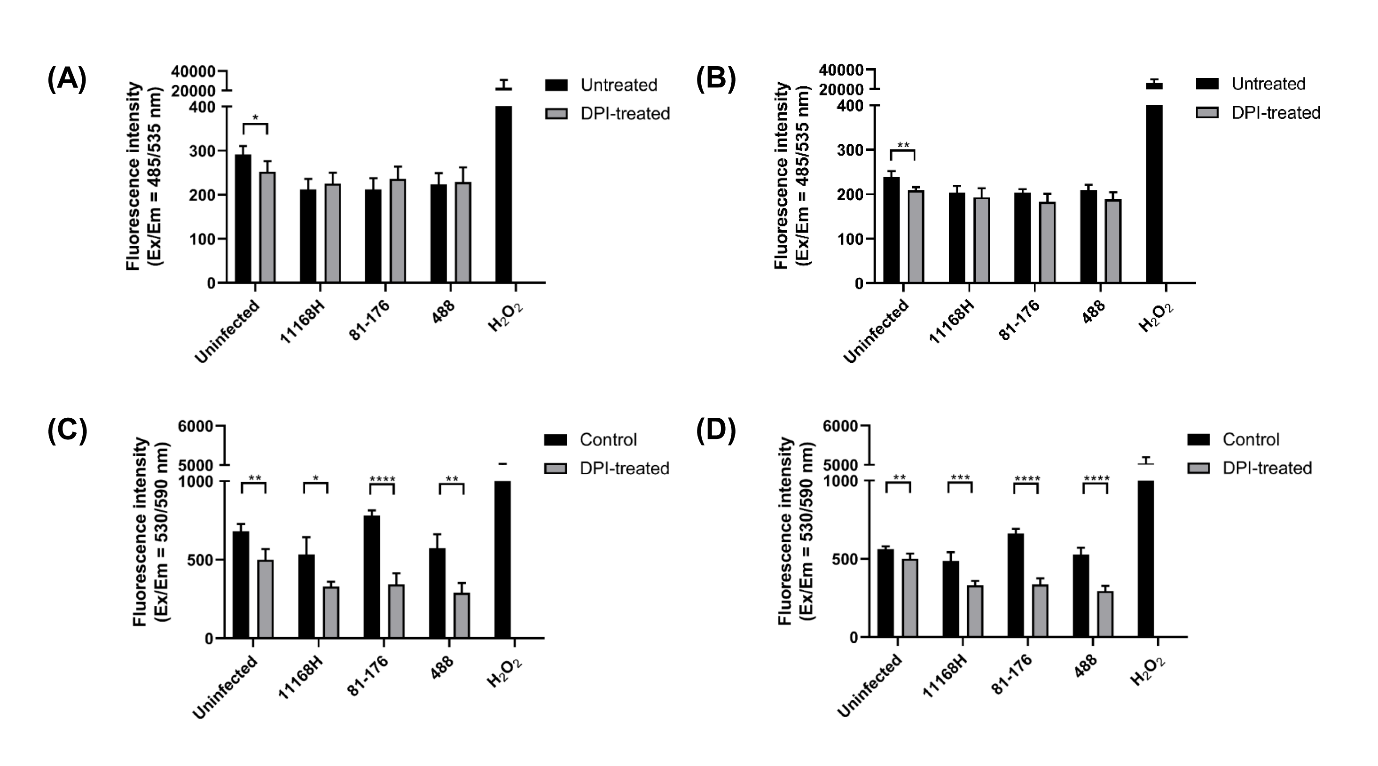

**FIGURE S1.** Detection of intracellular ROS and extracellular ROS in T84 and Caco-2 cells with or without DPI pre-treatment followed by C. jejuni infection. T84 and Caco-2 cells were pre-treated with 10 µM DPI for 1 hour and infected with C. jejuni. Intracellular ROS in (A) T84 and (B) Caco-2 cells after co-incubation with C. jejuni for 3 hours and extracellular ROS from (C) T84 cells and (D) Caco-2 after co-incubation with C. jejuni for 3 hours were measured. For detection of intracellular ROS, DCFDA was used. For detection of extracellular ROS, Amplex^®^ Red reagent with HRP were used. For positive controls, H_2_O_2_ was used. Experiments were repeated in triplicate. Asterisks denote a statistically significant difference (* = p < 0.05; ** = p < 0.01; *** = p < 0.001; **** = p < 0.0001).
