## Supplementary Figure S2 for "*Campylobacter jejuni* modulates reactive oxygen species production and NADPH oxidase 1 expression in human intestinal epithelial cells"

**Supplementary Figure 2**

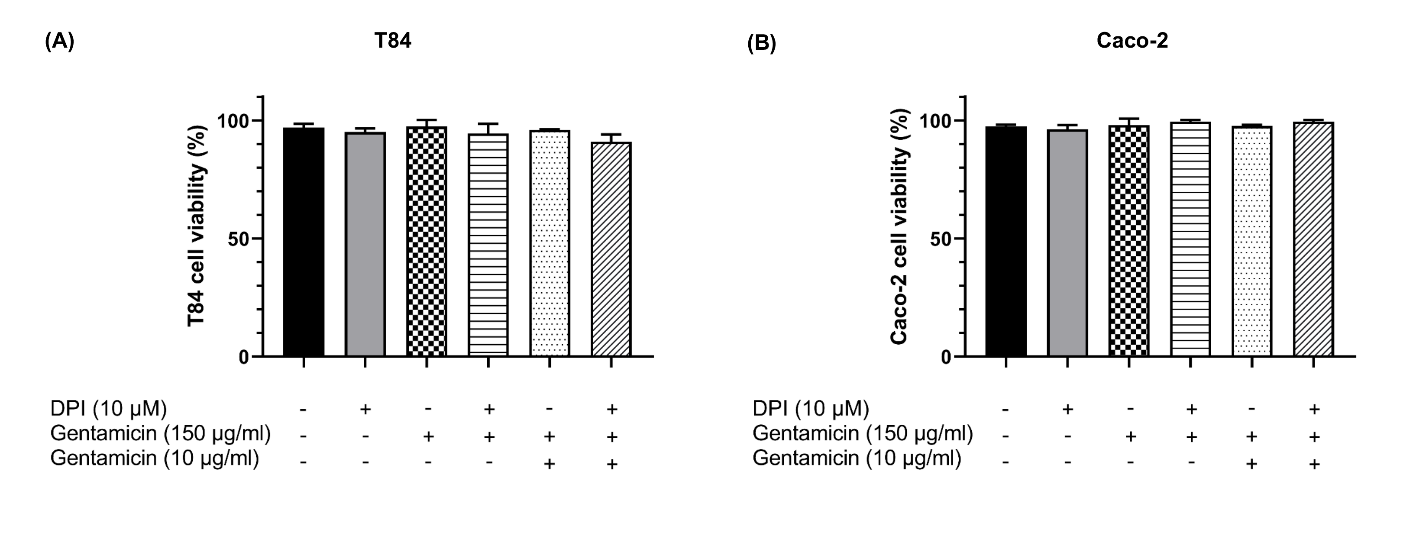

**FIGURE S2.** Trypan blue exclusion assay. (A) T84 and (B) Caco-2 cells were treated with DPI and stained with trypan blue dye to check their viability (%).
