## Supplementary Figure S3 for "*Campylobacter jejuni* modulates reactive oxygen species production and NADPH oxidase 1 expression in human intestinal epithelial cells"

**Supplementary Figure 3**

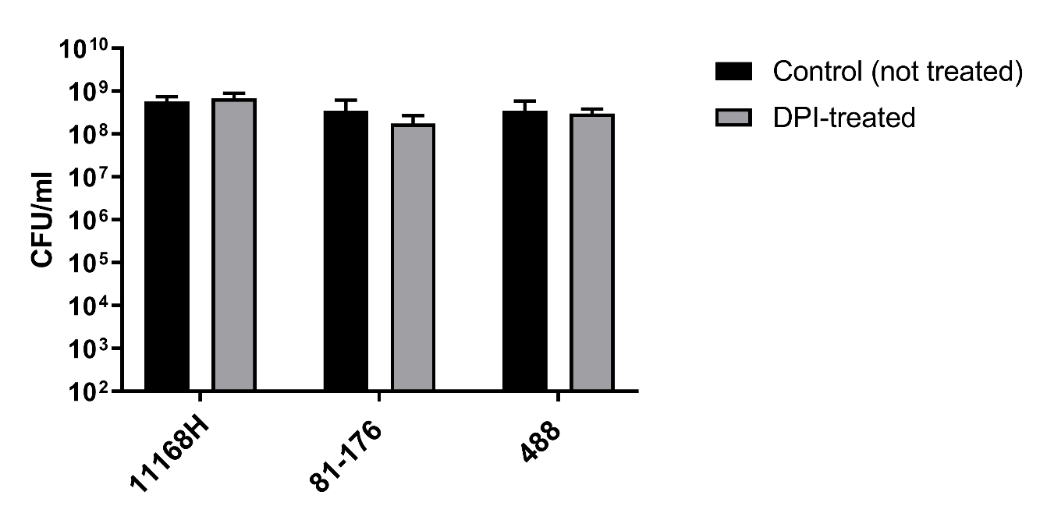

**FIGURE S3.***C. jejuni* viability assay with DPI treatment. T84 cells were treated with DPI for 1 hour and the cells were washed three times with PBS. After 1 hour DPI treatment, *C. jejuni* strains were co-incubated for 1 hour with PBS from the last wash and CFU/ml was recorded.
