## Supplementary Figure S4 for "*Campylobacter jejuni* modulates reactive oxygen species production and NADPH oxidase 1 expression in human intestinal epithelial cells"

**Supplementary Figure 4**

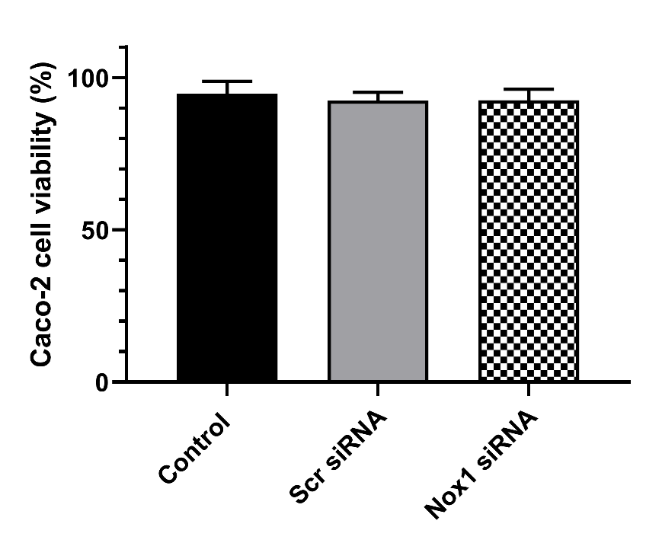

**FIGURE S4.** Trypan blue exclusion assay. Caco-2 cells were transfected with scrambled siRNA or Nox1 siRNA for 72 hours and stained with trypan blue dye to check their viability (%). Control is without any siRNA transfection.
