## Supplementary Table S1 for "*Campylobacter jejuni* modulates reactive oxygen species production and NADPH oxidase 1 expression in human intestinal epithelial cells"

**Supplementary Table 1**

**TABLE S1** Campylobacter jejuni strains used in this study

| *C. jejuni* strain | Description | Reference |
| --- | --- | --- |
| 11168H | A hypermotile derivative of NCTC 11168 wild-type strain which is a human clinical isolate. | (Karlyshev, Linton, Gregson, & Wren, 2002; Parkhill et al., 2000) |
| 81-176 | A human clinical isolate identified during an outbreak of acute enteritis associated with consumption of contaminated milk. | (Korlath, Osterholm, Judy, Forfang, & Robinson, 1985) |
| 488 | A human clinical isolate from Brazil which has a Type VI Secretion System (T6SS) | (Davies et al., 2019; Liaw et al., 2019) |
