## Supplementary Table S2 for "*Campylobacter jejuni* modulates reactive oxygen species production and NADPH oxidase 1 expression in human intestinal epithelial cells"

**Supplementary Table 2**

**TABLE S2** Primers used in this study

| Primer name | Sequences | qRT-PCR/ RT-PCR | Source |
| --- | --- | --- | --- |
| *nox1* | Forward: 5’-CACAAGAAAAATCCTTGGGTCAA-3’  Reverse: 5’-GACAGCAGATTGCGACACACA-3’ | qRT-PCR  and  RT-PCR | (Manea, Todirita, Raicu, & Manea, 2014) |
| *cat* | Forward: 5’-TGCAAGCTAGTGGCTTCAAAA-3’  Reverse: 5’-TCCAATCATCCGTCAAAACAA-3’ | qRT-PCR | (Wang & Eskiw, 2019) |
| *sod1* | Forward: 5’-GGCAAAGGTGGAAATGAAGAA-3’  Reverse: 5’-GGGCCTCAGACTACATCCAAG-3’ | qRT-PCR | (Wang & Eskiw, 2019) |
| *gapdh* | Hs_GAPDH_1_SG QuantiTect Primer Assay, QT00079247 | qRT-PCR | Qiagen |
| *gapdh* | Forward: 5’-CATCACCATCTTCCAGGAGC-3’  Reverse: 5’-GGATGATGTTCTGGAGAGCC-3’ | RT-PCR | (Das et al., 2000) |
